# Bulk-based hypothesis weighing increases power in single-cell differential expression analysis

**DOI:** 10.1101/2025.04.15.648932

**Authors:** Pierre-Luc Germain, Jiayi Wang, Mark D. Robinson

## Abstract

Due to the costs of single-cell sequencing, sample sizes are often relatively limited, sometimes leading to poorly reproducible results. In many contexts, however, larger bulk RNAseq data is available for the same conditions or experimental paradigm, which can be used as additional evidence of a generalizable differential expression pattern. Here, we show how such data can be used, via bulk-based hypothesis weighing (bbhw), to increase the power and robustness of single-cell differential state analysis. We find that all methods improve performance, with the best results obtained by applying a grouped Benjamini-Hochberg procedure on bins based on proportion-adjusted significance (PAS). These methods are implemented in the *muscat* package, and should be applicable to a broader range of scenarios.

## Introduction

The development of single-cell technologies has enabled the analysis of complex tissues at the resolution of individual cell types. However, due to the costs of these technologies, sample sizes are often relatively limited, sometimes leading to poorly reproducible results, especially with clinical data (e.g. Maleki et al., 2019; Nakatsuka et al., 2025). In many contexts, however, larger bulk RNAseq datasets are often available for the same conditions or experimental paradigm, which could be used as additional evidence of a differential expression pattern. Of course, many changes observed in specific cell types – especially rare ones – might not be visible in bulk data, and such a prior should not reduce our ability to detect novel changes. Here, we explore some simple methods to use bulk RNAseq data to increase the power and robustness of (multi-sample, multi-condition) single-cell differential expression analysis.

Consider as an example the recent data from Waag et al., 2025 on the response to stress in the hippocampus, where a large bulk RNAseq data (195 samples) is complemented by a smaller single-cell RNA-seq dataset. For each cell type, pseudobulk aggregation was performed to test for differential expression upon stress (Crowell et al., 2020), and the distribution of p-values (here for all neuronal cell types) is shown in Figure 1A. The p-value distribution shows a small excess of low p-values in comparison to the flat distribution that would be expected in the absence of genuine differences. While this is suggestive of a biological effect, the excess is very small, and consequently only a small fraction of genes can be reported as significant after False Discovery Rate (FDR) correction. However, we can partition the genes into quantile bins based on their significance in the much larger bulk dataset using the bulk RNA-seq data (Figure 1B). If we then look again at the distribution of single-cell p-values, but splitting genes according to this prior evidence (i.e. according to the significance bin in the bulk dataset), we can appreciate that some bins (e.g. bin 1) show a much larger excess of small p-values, while others (e.g. bin 3) are rather depleted of small p-values (Figure 1C). As a consequence of this greater (local) excess, the probability that any of those p-values is due to chance becomes lower, and thus a notable gain in power could be obtained by applying FDR to each bin separately.

**Figure 1.**
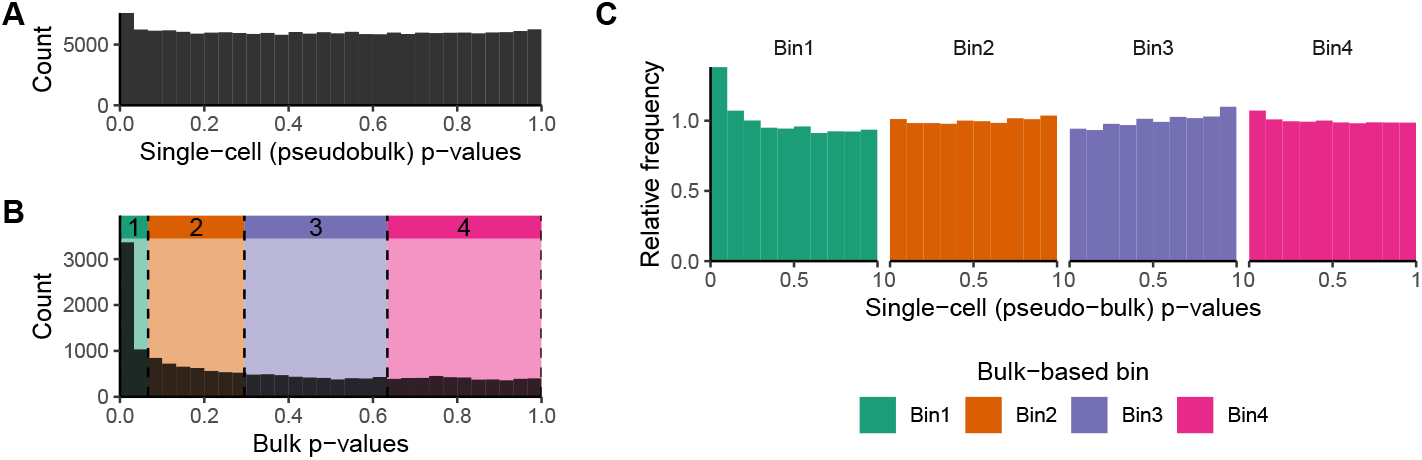
Example p-values of differential expression upon stress, from both a single-cell dataset with modest sample size (**A** and **C**) and a much larger bulk dataset (**B**) Waag et al., 2025. **C:** Splitting the single-cell p-values by significance of the respective gene in the bulk data leads to higher local excess of p-values.

Several methods have been proposed to harness such grouping of the hypotheses during FDR correction (Korthauer et al., 2019), in particular the Grouped Benjamini-Hochberg (gBH) procedures (Hu et al., 2010) and Independent Hypothesis Weighing (IHW) (Ignatiadis et al., 2016). gBH procedures work by estimating the proportion of true null hypotheses in each bin, which can be done through either the Least-Slope (LSL, Benjamini and Hochberg, 2000) or Two-Stage (TST, Benjamini et al., 2006) methods, and weighing the p-values accordingly (Hu et al., 2010). IHW instead uses a cross-validation strategy to optimize weights that lead to the highest rejections of the null hypothesis (see Ignatiadis et al., 2016).

In this brief report, we investigate the application of such adjusted hypothesis testing regimes, and additionally show that taking into consideration the local informativeness of the prior, i.e. the extent to which the cell type contributes to the bulk expression for each gene, further improves both precision and recall.

## Methods

### Example stress data

The example stress data used in Figures 1 and 3A was taken from the neuronal populations of (Waag et al., 2025). Specifically, we used the differential expression analysis results of dropping all acute stress related covariates from the model, i.e. Sex+CRS*TimePoint vs Sex+CRS (for the bulk, without Sex for the single-cell data that was only on females). For Figure 3A, we used a single cell type: ExN.CA1-do. Given the very strong differences in dynamics between the chiefly unprocessed RNA captured by single-nucleus RNAseq and standard bulk RNA, we used as a prior the minimum of the adjusted p-values computed on the full transcriptome or unspliced fraction of the reads.

### Toy p-value simulations

To explore the behavior of adjustment methods, we generated distributions of p-values (Figure 3B-D). For null simulations, we simulated 80k p-values using a uniform distribution. For simulations with a signal, we generated 74k uniform p-values (true null hypotheses), and complemented them with the absolute of normally-distributed values centered at 0, using 1000 with a standard deviation of 0.001, 3000 with 0.01, and 2000 with 0.025. Uninformative bins were generated by randomly sampling the desired number of bins with equal probability. Every scenario was simulated with 10 random seeds.

To generate informative bins, we first split the true null hypotheses (i.e. p-values generated with a uniform distribution) into 5 equally-size bins, and grouped the other p-values (i.e. with false null hypothesis) according to the p-value distribution from which they were sampled. We created noise by randomly assigning one third of the true null hypotheses to any of the bins with signal.

### scRNA-seq simulation

To evaluate different approaches, we first simulated two single-cell RNAseq data using muscat: one based on the reference dataset published with the package (Crowell et al., 2020) and the other based on a public dataset that profiled peripheral blood mononuclear cells (PBMCs) (Parse Biosciences, 2025). For the former dataset, we simulated two different groups (with 8 samples each) and six cell types (Figure 2A, C): three abundant cell types, only one of which differs between groups, and three rare, of which two differ between groups. The two affected rare cell types differ chiefly in that smallAffected1 has an expression profile similar to one of the large unaffected clusters, while smallAffected2 is more distinct but has fewer differentially-expressed genes (15% vs 20%, with an average logFC of 0.4). In this way, we simulate cases ranging from the ideal (an affected, abundant cell type), to the most challenging (a rare cell type with a very distinct response profile).

**Figure 2.**
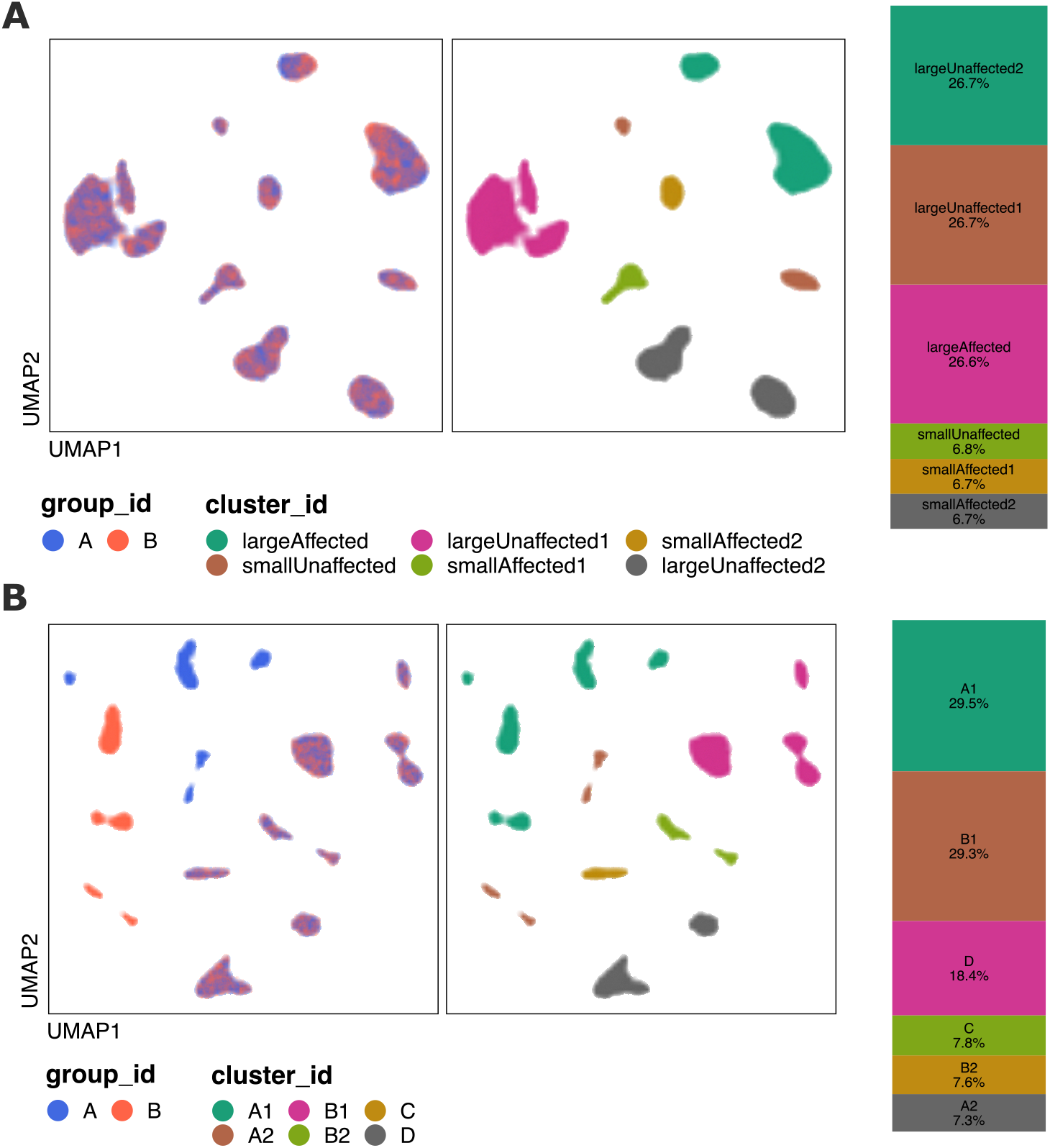
**A:** UMAP and cell type proportions for simulated dataset 1 (based on a mouse brain reference) with six cell types across two groups (8 samples each). **B:** UMAP and cell type proportions for simulated dataset 2 (based on human PBMCs) with six cell types across two groups (15 samples each).

**Figure 3.**
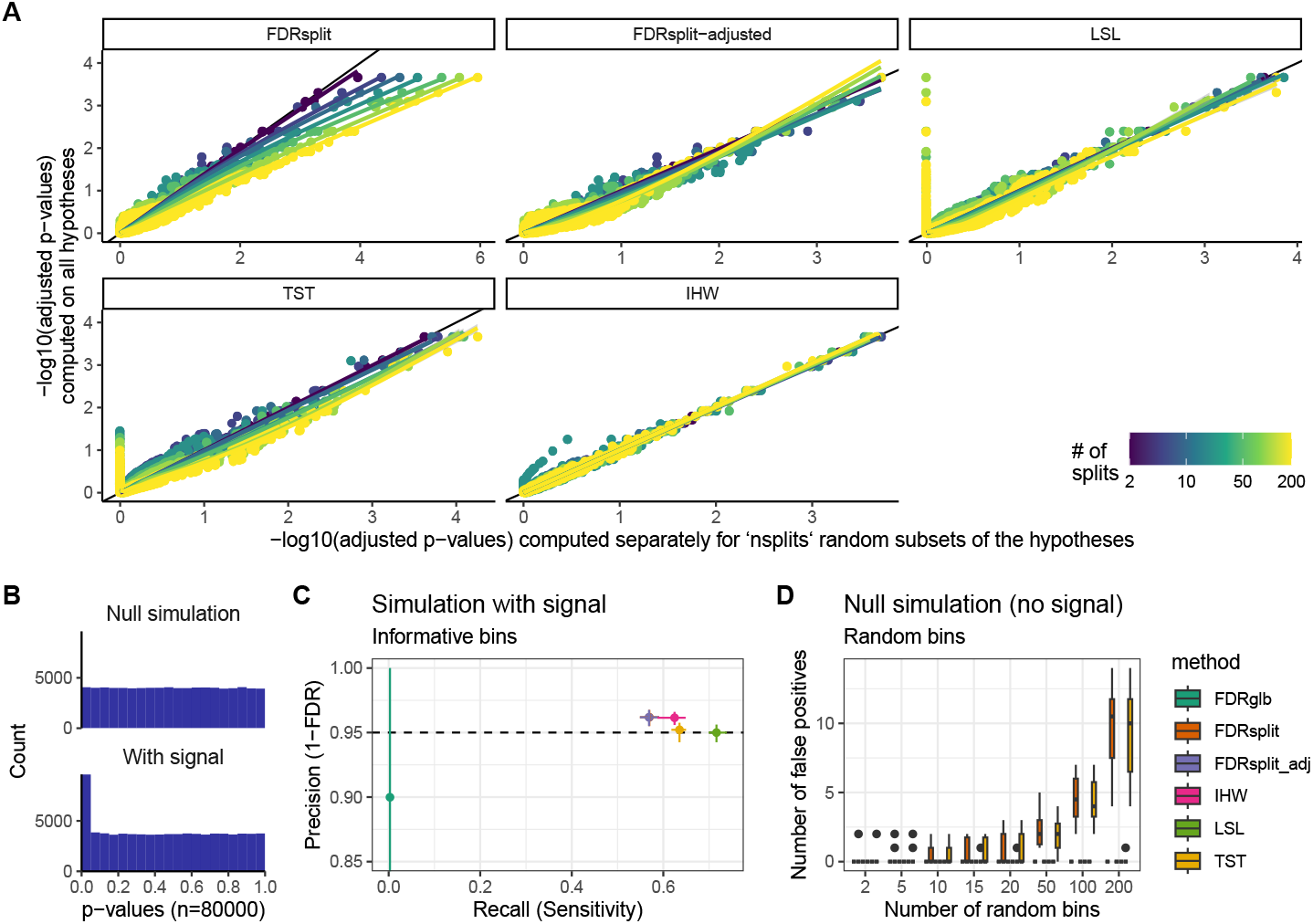
**A:** Adjusted p-values obtained by the different methods on the stress dataset with increasing numbers of uninformative (i.e. random) bins. **B:** Example toy simulations of p-value distributions with and without signal. **C:** Precision and recall (at nominal p.adj<0.05) of the adjustment methods in simulation with signals, using 8 informative bins. The dots indicate the average, and the segment the range, across 10 random seeds. (Note that the FDRsplit and FDRsplit-adjusted methods overlap here). **D:** Number of false positives reported by the methods in the null simulations with increasing number of un-informative (i.e. random) bins.

For the PBMC-based dataset, we simulated two groups of 15 samples each, again with a mixture of abundant and rare cell types (Figure 2B, C). Specifically, clusters A1/B1 (abundant) and A2/B2 (rare) represent large- and small-effect populations, respectively (large effect: 25% of genes with an average log2FC of 0.5; small effect: 25% of the genes with an average log2FC of 0.15). Cluster C (rare) exhibits a medium effect (20% of genes with an average log2FC of 0.3), while cluster D (abundant) does not differ between groups. Increasing the sample size to 15 per group in this setting allowed us to investigate how power changes with sample size.

For the evaluation, the 16 samples were summed to generate artificial bulk data, on which a differential expression analysis was performed using edgeR (Robinson et al., 2010), and 12 random sets of 2 vs 2 samples were used for pseudo-bulk differential state analysis using muscat-edgeR (Crowell et al., 2020).

### Downsampled real data

To have a complementary evaluation based on real data, we used the large single-nuclei dataset of human multiple sclerosis (MS) from Macnair et al., 2025, specifically restricting ourselves to white matter samples. We performed a pseudo-bulk differential analysis between the 13 control samples and the 51 diseased samples. As ground truth, we took the hypotheses with a (locally) adjusted p-value below 0.1 as true positives, and those above 0.5 as true negatives – the hypotheses in-between (24% of the genes) were judged unclear and discarded for the purpose of evaluation. We note that this definition of the negatives is a formally incorrect interpretation of the test, that is used as a necessarily inadequate proxy for a non-existing ground truth. Only cell types with more than 2 true positives were considered for the evaluation. We then tested the different methods on 12 random subsets of 4 control vs 5 diseased samples.

The MS data has strong condition-associated changes in cell type abundance, which would dominate bulk data. Therefore, to generate artificial bulk data, we first equalized the cell type abundance across samples, before summing the counts.

### Tested methods

We tested different ways of grouping hypotheses based on the bulk data, as well as different ways of using this grouping for p-value adjustment, all of which are now implemented in the bbhw function of the *muscat* package.

*Creation of the evidence bins*. We tested the following ways of grouping hypotheses based on the bulk data:

**p:** bins are based on quantiles of the bulk p-values. Missing (i.e. NA) p-values are put in their own bin if there are at least 50 of them, and otherwise set to 0.5.

**p+sign:** for each hypothesis (i.e. gene in a given cell type), if the direction of the change is different between bulk and the cell type, the bulk covariate is set to 0.7. Bins are then based on this modified covariate as for ‘p’.

**combined:** : first, significance-based bins are created in the same fashion as in ‘p+sign’. Each significance bin is then further split into genes for which the cell type contributes much to the bulk, and genes for which the cell type contributes little.

**PAS (Proportion-Adjusted Significance):** : for each hypothesis, the bulk significance is first adjusted for discordant directions as described for previous variants. Then the covariate is adjusted based on the cell type contribution to the bulk of that gene using *inv*.*logit*(*logit*(*ρ*)^∗^ *sqrt*(*c*)), where *p* and *c* are respectively the bulk p-value and the proportion of bulk reads contributed by the cell type). Bins are then based on the quantiles of this modified co-variate.

**PA-logFC (Proportion-Adjusted logFC):** : same as for *PAS*, except that shrunk bulk logFCs (from the edgeR::prefFC function) are used instead of the significance. The logFCs are converted to signed rank and normalized between -1 and 1, so that the same procedure as PAS can be applied.

**asNA:** : for hypotheses for which the cell type contributes little to the bulk profile, the covariate (i.e. bulk p-value) is set to NA, resulting in the hypothesis being placed in a separate bin. Non-NA values are then split into quantile bins.

In all cases, the number of bins is determined in relation to the number of hypotheses, ensuring a minimum of 1000 hypotheses per bin (except for the NA bin), and using a maximum of 12 bins except where indicated otherwise.

*p-value adjustment*. Once the bins are created, the following options were tested to make use of them:

**binwise:** : the Benjamini-Hochberg (BH) procedure is applied separately for each bin. Doing this can lead to an increase in false positives if the number of bins is large, and to correct for this the resulting adjusted p-values are multiplied by min(1,nbins*/*rank(*ρ*)), which results in proper FDR control even across a large number of bins.

**IHW:** : The Independent Hypothesis Weighing (IHW) method by Ignatiadis et al., 2016 is applied, using the *IHW* R package with the default alpha of 0.1. If the 25% quantile of the bin sizes was above 1000, 5-fold validation was performed. If it was above 400, 4-fold, and otherwise 3-fold.

**gBH.LSL:** : Grouped Benjamini-Hochberg (gBH) is used (Hu et al., 2010) with the Least-Slope (LSL) estimation of the proportion of true null hypotheses (Benjamini and Hochberg, 2000).

**gBH.TST:** : Grouped Benjamini-Hochberg (gBH) is used (Hu et al., 2010) with the Two-Stage (TST) estimation of the proportion of true null hypotheses. The Two-Stage approach amounts to using BH within each group, and use the rejection rate at a given nominal FDR threshold (here we used 0.05) to estimate the proportion of true null hypotheses within the group.

Each method was run in two flavors: a local one, which is applied for each cell type separately, and a global one, which is applied once across all cell types.

## Results

### General behavior of the adjustment methods

We first tested the different methods that leverage hypothesis grouping using the Waag et al. (2025) stress single-cell data. While there is no ground truth in this data, we tested the expectation that using uninformative (i.e. random) bins should not lead to systematically higher reported significance. As shown in Figure 3A, computing adjusted p-values (using the FDR method) separately for the different bins (FDRsplit) leads to a systematic decrease in the reported adjusted p-values, although the effect was very mild unless the number of bins was very high. This could be easily corrected by multiplying the adjusted p-values by max (1 *n/*rank (*padj*)), where *n* is the number of bins and rank (*padj*) is the rank of the (pooled) adjusted p-values. The results are shown as ‘FDRsplit-adjusted’. The Two-Stage grouped BH method (TST) had a similar trend, but much milder, while the other methods did not lead to a systematic FDR underestimation.

### All methods leveraging bulk-based evidence bins improve power

We next turned to our specific case of using bulk data to improve power of per-cell-type differential expression across condition in single-cell data. To this end, we used simulated data, as well as downsampled real data with the full-sized data serving as ground truth (see Methods). All methods led to an improvement in the precision-recall curves, mostly through an improvement in recall, except for the adjusted bin-wise method that instead led to increased precision (Figure 4A-B). The two grouped BH variants (LSL and TST) showed the strongest improvement.

**Figure 4.**
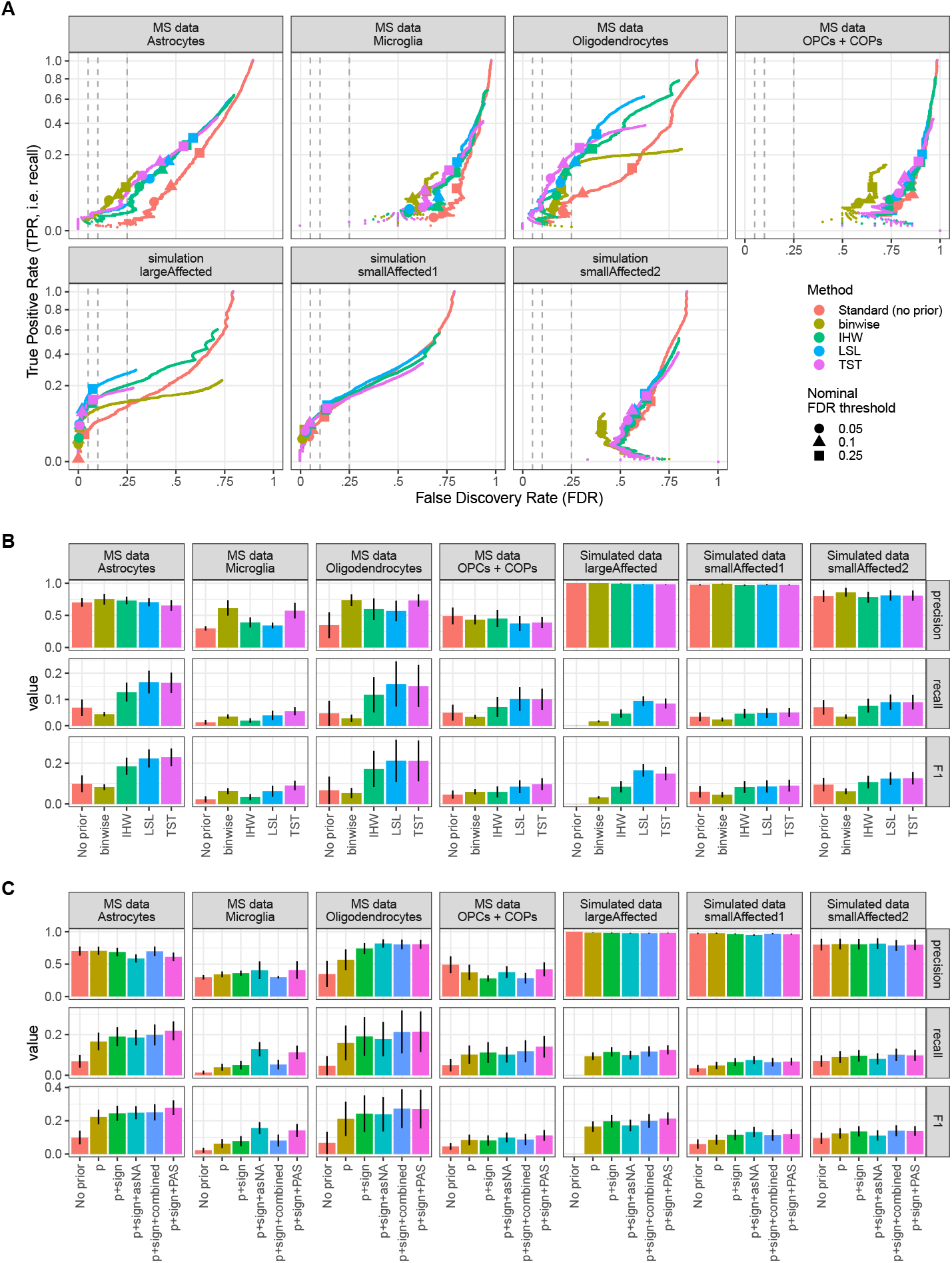
**A:** TPR-FDR curves of the different adjustment methods, using bins based on quantiles of raw bulk significance. The dashed lines indicate an effective FDR of 0.05, 0.1 and 0.25. **B:** Precision, recall and F1 score of the same methods at a nominal adjusted p-value threshold of 0.05. **C:** Precision, recall and F1 score of the top method (gBH-LSL) using different ways to group the hypotheses (i.e. variants of the bin preparation). ‘p’ stands for defining bins based on the bulk p-value only. ‘p+sign’ stands for discounting opposite directions. The other three methods additionally take the proportion of contribution into account (see Methods). In **B-C**, the error bars indicate the standard error of the mean across random seeds (note that most of the variation is due to baseline difference in power between the pseudobulk samples selected).

### Further improvements by considering direction and cell type contribution to the bulk

The prior evidence, i.e. the bulk data, is not equally informative for all genes and cell types. It is more informative for abundant cell types than rare cell types, because these influence the bulk more, and it is also more informative for a gene that is chiefly expressed in the cell type of interest, versus a gene that is mostly expressed in other cell types (which would dilute, at the bulk level, the effects in the cell type of interest). We therefore computed, based on the pseudobulk profiles, the proportion of the bulk reads expected to be attributable to each cell type for each gene, and tested ways to include that in the creation of the evidence bins (see Methods). In addition, we discounted (i.e. increased) the bulk p-values when the bulk logFC was in the opposite direction to that of the cell type of interest. To test these changes, we first focused on the top adjustment method (gBH-LSL) (Figure 4C). Taking the direction of change into account led to a systematic improvement. However, not all bin construction methods using the proportion of contribution into account had positive impact, but the Proportion-Adjusted Significance (PAS) and combined approaches generally brought further slight improvements (Figure 4C). Repeating this for all combinations of binning and adjustment methods (Figure 5A) confirmed the robustness of this approach across adjustment methods.

**Figure 5.**
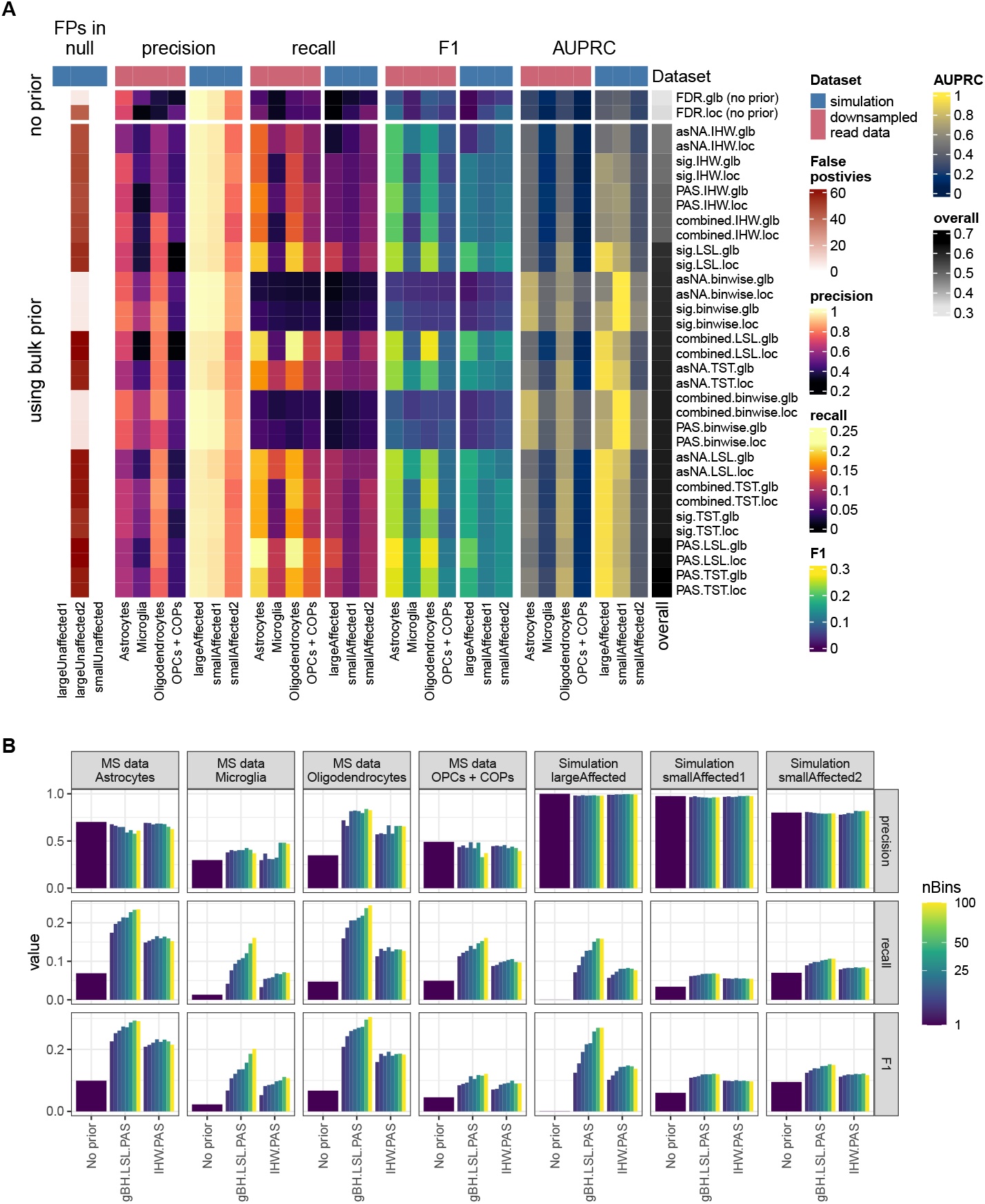
**A:** Precision, recall, F1 score, area under the precision-recall curve (AUPRC), and (in unaffected cell/types) number of false positives reported by each combination of bin- and adjustment-method. **B:** Precision, recall and F1 scores when using different numbers of bins. All values are at a nominal adjusted p-value threshold of 0.05.

### Effect of the number of bins used

We tested the impact of the number of bins used on the results (Figure 5B). Both IHW and gBH were fairly robust to the number of bins. There was an initial gain in recall for all methods when increasing the number of bins from 5 to 8 and 12, but beyond this IHW did not show any improvement. This is presumably because the cross-validation strategy, which requires a certain number of hypotheses per bin to work properly. Instead, gBH sometimes showed further increase in recall with larger numbers of bins, although the optimum differed across cell types.

### Similar results using shrunk logFC instead of significance

It has often been argued that p-values are not the best way of ranking genes, and that fold-changes can be more appropriate, although involving its own risks (Xiao et al., 2012). As the accuracy of fold-change estimates in sequencing data is notoriously dependent on that of the quantification, we tested whether using bulk-level, shrunk logFC estimates (Phipson, 2013) could provide a better grouping of the hypotheses. While both approaches were superior in some contexts, overall they appeared to provide a very similar increase in power (Figure 6).

**Figure 6.**
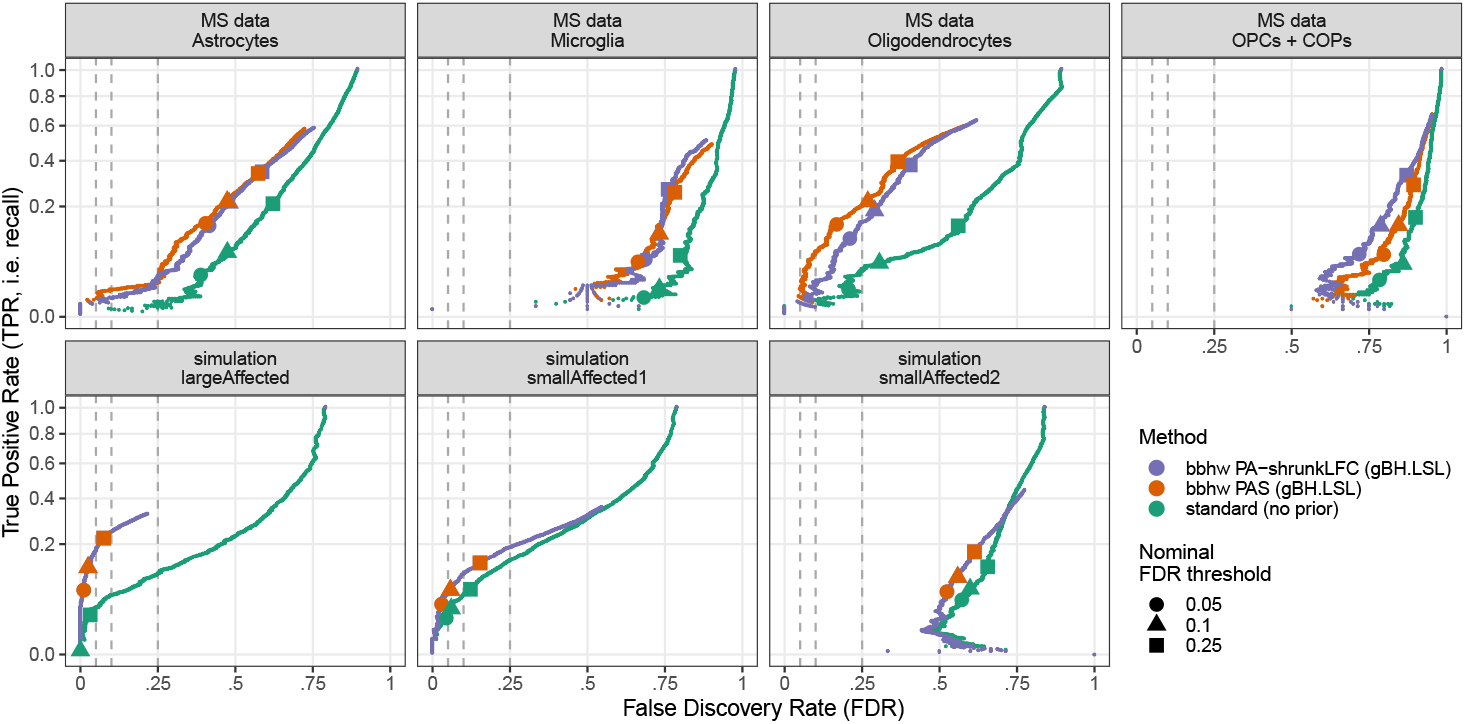
TPR-FDR curves of the top method based either on bulk significance or shrunk logFC, compared to standard (no-prior) FDR.

### Effect of sample size in an independent simulation

We hypothesized that the gains from bulk-based hypothesis weighing are most likely strongest when the single-cell data has a small sample size, as in the examples above (2vs2 for the simulated mouse data, and 4vs5 for the real human data). To investigate this, as well as confirm the previous findings, we developed a new simulation based on a different reference dataset, derived from a different technology: the 12 healthy donors from the Parse Biosciences 1 million PBMCs dataset (Parse Biosciences, 2025). This second simulation largely confirmed the previous results, this time with combined-LSL showing the best overall performance along with PAS-LSL (Figure 7A). To our surprise, the gain in power was not restricted to low sample sizes, and substantial improvements were still observed when the single-cell data was better powered (Figure 7B).

**Figure 7.**
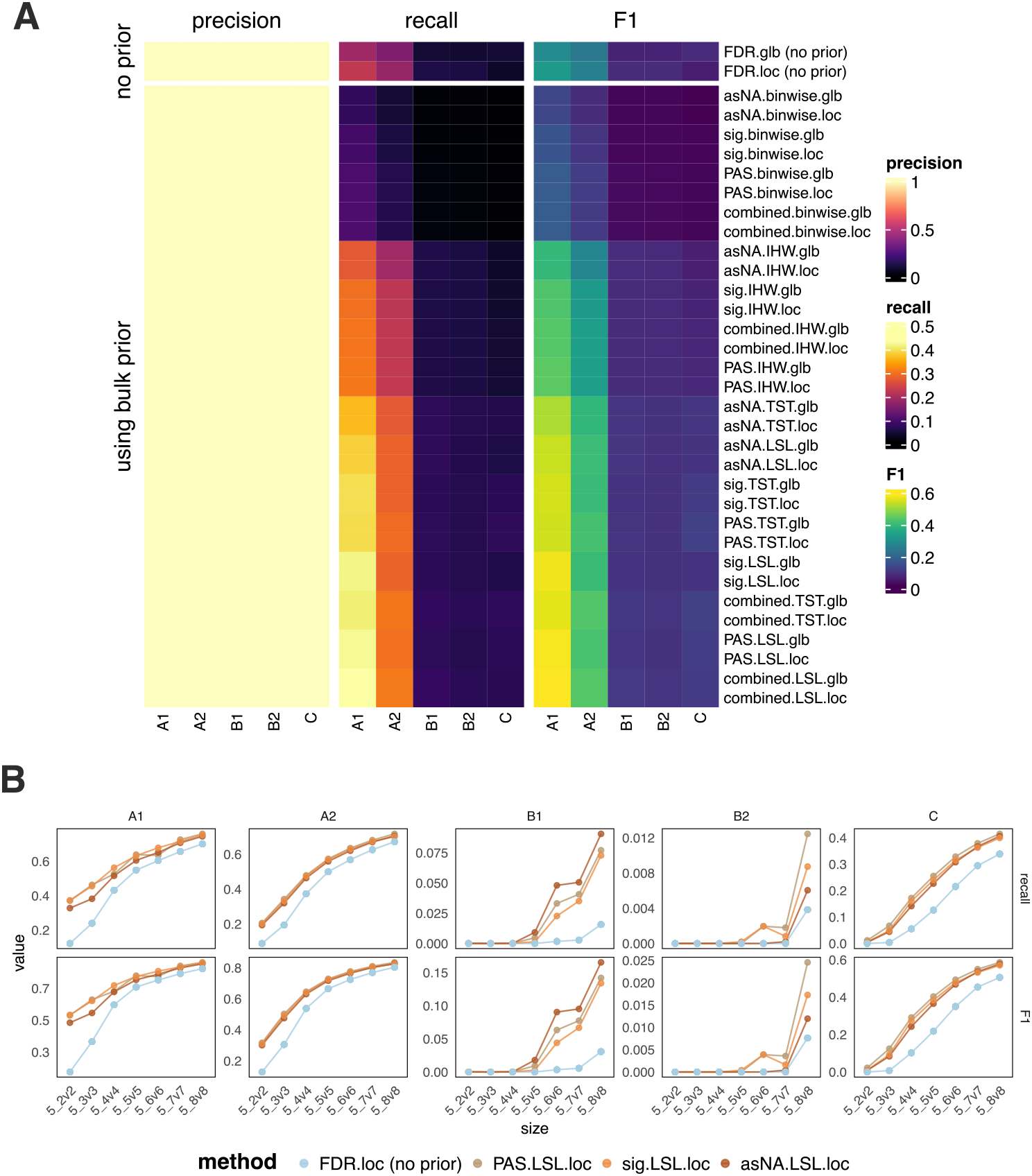
**A:** Precision, recall, F1 score reported by each combination of bin- and adjustment-method. **B:** Recall and F1 changes with different sample sizes used in single-cell data.

### Illustration on the stress dataset

Finally, we illustrate the application of bbhw on the dataset from Waag et al., 2025 on the hippocampal response to acute stress, previously discussed in Figure 1. This represents a real-life scenario, using real bulk RNA-seq with important technological differences to single-nuclei RNAseq data, and timepoints that are not exactly matched between the two Waag et al., 2025 datasets. Given these differences, we used as a prior the minimum of the adjusted p-values computed on the full transcriptome or unspliced fraction of the reads. Despite these differences, bbhw lead to a major increase in power (Figure 8. The gained DEGs were very strongly enriched (12.5-fold enrichment, Fisher’s exact test p<2.2e-16) for primary neuronal response genes as defined by Tyssowski et al., 2018, as well as for genes responding to activation of the glucocorticoid receptor (16-fold enrichment, p=1e-10) from Gerstner et al., 2022, supporting to their biological relevance.

**Figure 8.**
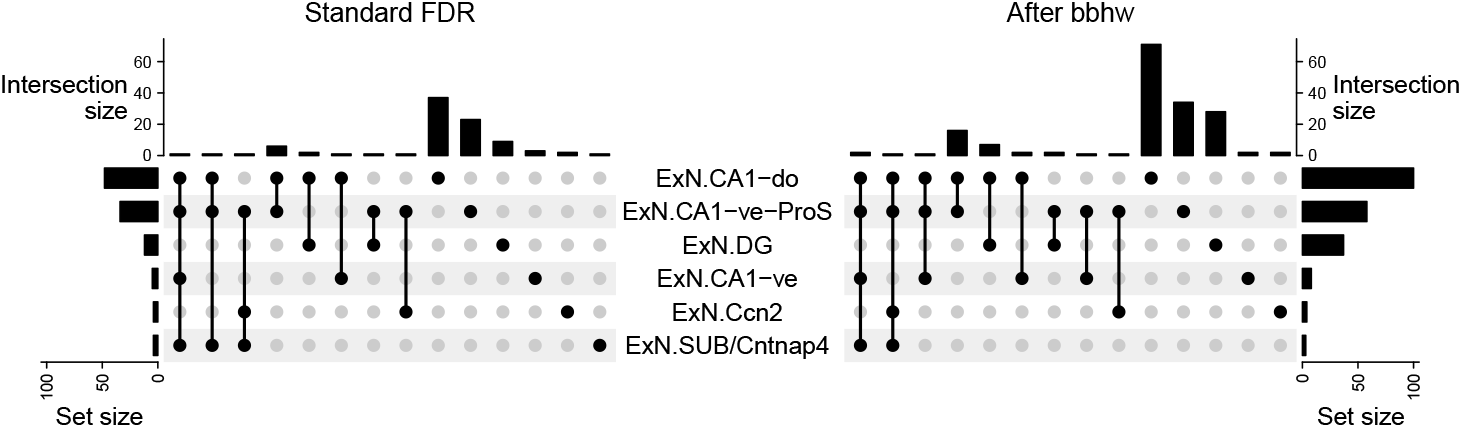
UpSet plot showing the number and overlap of the DEGs identified using standard FDR correction (left) and bbhw (right). Shown are the main responding neuronal cell types.

## Discussion

We have shown how simple strategies based on grouped FDR correction can leverage bulk data in order to increase the precision and/or sensitivity of single-cell differential state analysis. The magnitude of gain in power can be expected to strongly vary across datasets, and across cell-types within these, especially depending on their abundance and/or similarity to abundant cell-types. The methods discussed here are now all implemented in the *muscat* bioconductor package. Because they operate at the level of multiple correction, they can however be used in combination with any differential analysis package. The methods could also be directly applied to other problems, such as eQTL discovery in single-cell data (Bryois et al., 2022).

While the methods had good FDR control in the simulations, FDR was generally underestimated in the real data. We believe that there are two main reasons for this. First, in order to establish a ground truth, we removed from the evaluation a lot of hypotheses that were considered ambiguous (i.e. those which had an adjusted p-value between 0.5 and 0.1 in the full data), which means that we are not computing precision on the full set of rejected hypotheses. This is a limitation of this study, for which the accompanying simulations offers some mitigation. Second, the ground truth established from the larger cohort effectively tests the generalizability of the findings, and given patient heterogeneity, it is very likely that many differences are true of the smaller comparison (i.e. would be reproduced if other samples of the same individuals had been taken) but not generalizable across the population.

In the case of the MS data, we had to standardize the cell type abundance of the ‘artificial’ bulk data in order for the bbhw-methods to genuinely improve performance, otherwise the bulk profiles would be dominated by differences in cell type proportions. Obviously, this cannot be done with real bulk data. Therefore, a limitation of bulk-based hypothesis weighing is that it is unlikely to yield much improvement when the experimental condition studied involves major expansion or depletion of some cell types. However, in such cases, we also do not expect the method to deteriorate the results, making it safe for application.

How informative the bulk data is for the single-cell differential analysis depends on how closely aligned the experimental designs were, but is also influenced by technological differences between the two. For example, bulk RNAseq typically covers the entire gene, increasing sensitivity for larger genes, while the most common single-cell technologies chiefly cover the 3’ end. Such differences, which are not recapitulated in the datasets used for benchmark here (for which the bulk data is obtained by aggregating single-cell data), constrain the power that can be gained from bbhw. Similarly, while most transcripts captured by single-nuclei data are unprocessed, this is a small minority in standard bulk data, and in such circumstances we recommend using the unspliced fraction of the bulk data for hypothesis grouping/weighing. We illustrated this with the stress data, observing substantial gains in power despite technological differences.

Finally, an interesting finding of the study is that grouped Benjamini-Hochberg (Hu et al., 2010) (in particular using the Least-Slope estimator) outperformed Independent Hypothesis Weighing (IHW) (Ignatiadis et al., 2016), although the difference was small. This is in contrast with the results obtained by Ignatiadis et al. (2016), who found gBH ‘slightly anti-conservative’. If gBH underestimates FDR, it appears in our setting to be by very little. gBH also has the benefit of being deterministic and very fast to compute (we provide an implementation in the *muscat* package). While this might not generalize to other applications, both methods would be worth further study.

## Data, script, code, and supplementary information availability

The implementation of the methods can be found in the *muscat* package. The scripts to reproduce the analyses and figures can be found at https://github.com/plger/bbhw. An archived copy of both, with the specific versions used, can be found at https://dx.doi.org/10.5281/zenodo.17376376.

